# Emergent sub-population behavior uncovered with a community dynamic metabolic model of *Escherichia coli* diauxic growth

**DOI:** 10.1101/291492

**Authors:** Antonella Succurro, Daniel Segrè, Oliver Ebenhöh

## Abstract

Microbes have adapted to greatly variable environments in order to survive both short-term perturbations and permanent changes. A classical, yet still actively studied example of adaptation to dynamic environments is the diauxic shift of *Escherichia coli*, in which cells grow on glucose until its exhaustion, and then transition to using previously secreted acetate. Here we tested different hypotheses concerning the nature of this transition by using dynamic metabolic modeling. Towards this goal, we developed an open source modeling framework integrating dynamic models (ordinary differential equation systems) with structural models (metabolic networks), which can take into account the behavior of multiple sub-populations, and smooth flux transitions between different time points. We used this framework to model the diauxic shift, first with a single *E. coli* model whose metabolic state represents the overall population average, and then with a community of two sub-populations each growing exclusively on one carbon source (glucose or acetate). After introducing an environment-dependent transition function that determines the balance between different sub-populations, our model generates predictions that are in strong agreement with published data. We thus support recent experimental evidence that, rather than a coordinated metabolic shift, diauxie would be the emergent pattern of individual cells differentiating for optimal growth on different substrates. This work offers a new perspective on the use of dynamic metabolic modeling to investigate population heterogeneity dynamics. The proposed approach can easily be applied to other biological systems composed of metabolically distinct, interconverting sub-populations, and could be extended to include single-cell level stochasticity.

**Importance:** *Escherichia coli* diauxie is a fundamental example of metabolic adaptation that is not yet completely understood. Further insight into this process can be achieved by integrating experimental and computational modeling methods. We present a dynamic metabolic modeling approach that captures diauxie as an emergent property of sub-population dynamics in *E. coli* monocultures. Without fine tuning the parameters of the *E. coli* core metabolic model, we achieve good agreement with published data. Our results suggest that single-organism metabolic models can only approximate the average metabolic state of a population, therefore offering a new perspective on the use of such modeling approaches. The open source modeling framework we provide can be applied to model general sub-population systems in more complex environments, and can be extended to include single-cell level stochasticity.

## 1 Introduction

In natural environments microorganisms are exposed to high fluctuations of nutrient and micronutrient availability and have therefore evolved adaptation strategies, both short-term to respond to temporary perturbations and long-term to increase evolutionary fitness [1]. We still lack a sound theoretical understanding of the mechanisms driving such strategies, but the recent technological advances in high-throughput experimental techniques pave the way to novel approaches that integrate experimental and theoretical biology [2]. Theoretical ecology describes ecosystems in mathematical terms as dynamic organism-environment interactions [3]. As in statistical physics, individual behaviors in an *ensemble* result in observable emergent patterns that can be modeled with mathematical equations [4]. This is the case for the earliest models of population dynamics developed by Verhulst [5], Lotka [6] and Volterra [7] and for the pioneering work of Jaques Monod in modeling microbial growth [8]. With the rising academic and industrial interest in the “microbiome”, systems biology approaches are becoming a new standard [9] and more methods for the mathematical modeling of microbial communities are being developed [10, 11].

In constraint-based stoichiometric modeling the metabolic network model of an organism is reconstructed from its annotated genome and described mathematically as a stoichiometric matrix **S**. After imposing the steady-state assumption and introducing thermodynamic and biological boundaries for the metabolic fluxes 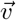, flux balance analysis (FBA) [12, 13] defines an optimization problem in order to identify one particular flux distribution in the solution space. As long as the objective function (which imposes further biological assumptions on the system) is linear in the fluxes, the optimization problem can be solved by linear programming (LP). FBA returns a unique solution for the objective function, but the metabolic flux distribution is generally not unique, especially in genome-scale metabolic network models (GEMs). Based on the hypothesis that metabolism has evolved to make efficient use of resources and minimize waste, two specific methods were developed to extend FBA: parsimonious FBA (pFBA) [14] and minimization of metabolic adjustment (MOMA) [15]. In pFBA a second LP is defined such that the value of the objective function is set to the FBA solution and the new objective is the minimization of the overall fluxes. MOMA was developed to simulate the response to the perturbation introduced by gene deletion and is based on the principle that the organism will readjust its metabolism to a minimally different configuration with respect to the wild-type optimum. Another extension of FBA, dynamic FBA (dFBA) [16], allows partial recovery of the dynamic information lost under the steady-state assumption. In the static optimization approach (SOA) that underlies dFBA, time is divided into discrete intervals and a new FBA problem is solved at time *i* after updating the external conditions according to the FBA solution at time *i* − 1. Approaches to model microbial communities with GEMs have been recently reviewed by Succurro and Ebenhöh [17].

FBA and dFBA have been applied to study one of the most basic examples of metabolic transitions: diauxie [16, 18, 19]. Discovered in the model organism *Escherichia coli* in 1941 by Monod [8, 20], diauxie remains a topic of active research [21, 22, 23]. Under aerobic conditions with glucose as the sole carbon source (also generally the preferred one), *E. coli* secretes acetate during growth, which it then consumes once the glucose is exhausted. The molecular mechanisms driving this transition are still not completely understood, but over the last few years the fundamental role of stochasticity and population heterogeneity has been demonstrated experimentally [24], often with the support of mathematical models. Indeed, in unpredictable natural environments with fluctuating nutrient availability and variable fitness landscapes, homogeneous populations are more likely to face extinction, and bet hedging provides a selective advantage [25]. Single-cell studies have suggested that the observed biphasic growth is possibly the effect of stochastic gene expression [21], eventually co-regulated by memory mechanisms [26]. Kotte *et al.* [27] systematically investigated bistability in a clonal *E. coli* population. After ruling out responsive switching as a homogeneous adaptation, their results strongly suggested that the heterogeneous adaptation that results in two co-existing phenotypes was driven by responsive diversification (with a single phenotype diversifying in response to environmental changes) rather than stochastic switching (where the two phenotypes would co-exist from the beginning). Although stochastic mathematical models have been proposed to support those findings, metabolic modeling approaches are only considered suitable to describe homogeneous systems, with single organism GEMs representing the average population metabolic state.

Varma and Palsson [18] performed the first dFBA analysis on *E. coli*, with a single GEM growing aerobically first on glucose and then on the secreted acetate. Here we present a study of *E. coli* diauxic growth on these two carbon sources, with the bacterial population modeled either as having an average, unique metabolic state (standard FBA and dFBA approach) or as the combination of two *E. coli* populations adapted to one of the two carbon sources. We use a modeling approach that integrates ordinary differential equation (ODE) models with dFBA, extending methods typically applied to study the dynamics of multi-species communities to the investigation of emergent patterns from individual behavior in monocultures. We implement three approaches: (i) we model a homogeneous yet smooth shift, with a single *E. coli* GEM, by adapting the MOMA algorithm; (ii) we introduce the hypothesis of sub-populations growing on specific carbon sources and model population transition as a purely stochastic mechanism; and (iii) we introduce an environment-driven response. Our results suggest that, rather than as a coordinated metabolic shift, diauxie can be modeled as the emergent pattern resulting from sub-populations optimizing growth on different substrates in response to environmental changes. This is much in agreement with experimental evidence from *e.g.* Kotte *et al.* [27], and offers a new perspective on the use of dynamic metabolic modeling to investigate population dynamics. The proposed approach can easily be transferred to studies of generic sub-populations or communities, and ultimately can be expanded to investigate single-cell dynamics.

## 2 Results

We ran simulations with an open source modeling framework developed to model ecosystem dynamics. The models are ODE systems solved with integrating routines that at each integration step solve an FBA problem. We first validated the *E. coli* GEM on the data from Varma and Palsson [18] (who reported the first dFBA analysis of the glucose-acetate shift) and then used the calibrated model to reproduce the independent sets of experiments from Enjalbert *et al.* [22] (who analyzed *E. coli* grown in aerobic batch systems with different concentrations of glucose and acetate). In the standard dFBA approach, a population is modeled with a unique GEM and fluxes instantaneously change to adapt to new environmental conditions. In reality, however, transcriptional changes and flux rerouting may cause delays, which are not captured by existing algorithms. Furthermore, dFBA might predict metabolic states in which more carbon sources are simultaneously utilized, and it is not obvious that such an approach will correctly capture the complexity of a population diversifying into metabolically distinct sub-populations. Therefore, we modified the dFBA algorithm taking advantage of optimization strategies previously developed for different biological questions and implemented novel concepts as well. In particular, we used either pFBA [14] or an adaptation of MOMA [15] to solve the FBA problem at each time step, replicating the standard dFBA approach or implementing a homogeneous yet smooth shift, respectively. The MOMA algorithm was integrated into the dFBA routine by imposing that the solution of the FBA problem at time *i* is minimally different from the solution at time *i* − 1. We tested three different hypotheses: (i) homogeneous, smooth population shift; (ii) stochastic-driven and (iii) environment-driven sub-population differentiation. We observed that dFBA with both pFBA and MOMA predicted abrupt transitions from acetate catabolism to acetate anabolism, and condition-specific parameterizations were necessary to reproduce the different data. We then modeled two *E. coli* sub-populations growing exclusively on glucose or acetate. For this we extended the standard dFBA approach to include the process of population shifts. We tested whether purely stochastic switches (ii) or rather a responsive diversification (iii) could capture the diauxic behavior by modeling the population transitions either with constant rates (ii) or with a heuristic function dependent on carbon source concentrations (iii). We observed that only model (iii) could reproduce data from different experiments with a unique set of parameters. We did not find significant improvements using MOMA rather than pFBA within the same metabolic state, so the simpler pFBA implementation was used in the sub-populations simulations where each model is fixed into one metabolic configuration. Further details of the modeling approach are provided in Materials and Methods.

### *E. coli* diauxie modeled with a uniform population

A single GEM was used to model the average *E. coli* metabolic state and we compared the simulation results with the original data from Varma and Palsson [18] (Fig. S1). The parameters for the simulations are reported in Tab. 1 and 2, and the only flux constraints that we calibrated to the data were the oxygen uptake rate and the maximal acetate secretion rate. A fixed cell death rate was introduced as previously described in [19], with a value from the literature (Tab. 1). In these simulations, a lower absolute flux variation at each simulation time step was observed with the MOMA implementation (Fig. S2). We used the same GEM model to reproduce the results from Enjalbert *et al.* [22], changing only the initial values of biomass, glucose and acetate (Fig. 1 and S5). Although the pFBA simulation (Fig. 1(a)) showed a brief shift to growth on acetate at the time of glucose exhaustion (~ 4 h), the MOMA simulation predicted complete growth arrest already at that point, with a minimal acetate consumption to satisfy the ATP maintenance requirement implemented in the GEM (Fig. 1(b)). Both simulations well captured glucose consumption and acetate secretion, but neither of them was able to reproduce the slow acetate consumption observed experimentally. Even after fine-tuning the constraint on acetate up-take to achieve a perfect match of the acetate consumption data from Varma and Palsson [18], the model could not reproduce the acetate concentration dynamics of the corresponding data from Enjalbert *et al.* [22] (data not shown). Therefore, we decided to avoid fine-tuning of acetate uptake (Tab. 1). Both pFBA and MOMA simulations showed an abrupt change in the flux distribution upon shifting from glucose to acetate consumption (Fig. S3). We evaluated the agreement between experiment and simulation with the *R*^2^ distance between *in vivo* and *in silico* data for the biomass (pFBA *R*^2^ = 0.989; MOMA *R*^2^ = 0.982) glucose (pFBA *R*^2^ = 0.993; MOMA *R*^2^ = 0.993) and acetate (pFBA *R*^2^ = 0.277; MOMA *R*^2^ = 0.409). In Fig. 2 we compare the flux distributions of our simulation results to the experimental results reported by Enjalbert *et al.* [22] for over/under-expression of key genes associated with glucose and acetate metabolism (represented graphically in the top panels). First, we computed the flux solutions for *E. coli* growing on either glucose or acetate exponentially (data not shown) and compared the fluxes through the relevant reactions in *E. coli* growing on acetate to those in *E. coli* growing on glucose. Fig. 2(a) shows the absolute values for the flux results in the two simulations, normalized between 0 and 1 for direct comparison with the qualitative representation of the gene expression data (with 0 for non-expressed and 1 for expressed genes, respectively). The simulation results were consistent with the experiments, with active reactions (dark green) related to acetate consumption and anabolism (ACKr, PPCK, FBP, ICL, MALS) and inactive reactions (white) related to glycolysis (PFK, PYK) during growth on acetate, and vice versa during growth on glucose. PPS did not carry flux in either simulation. We then used the simulation results presented in Fig. 1 to compare the metabolic fluxes before and after glucose exhaustion (GE), *i.e.* before and after the single *E. coli* model shifts from growth on glucose to growth on acetate. Enjalbert *et al.* [22] compared gene expression levels between samples taken at time (GE + 30 min) and (GE - 115 min). However Fig. 1 shows that according to the simulation, growth has already stopped after 30 min from the GE point. Indeed comparing the absolute values of fluxes taken at time (GE + 30 min) and (GE - 115 min), we found that both pFBA and MOMA simulations qualitatively captured the down regulation trends, whereas neither reproduced the observed up-regulation (data not shown). Fig. 2(b) shows the difference in absolute values of fluxes taken at time (GE + 18 min) and (GE - 115 min), where in pFBA simulations growth is still observed. In this case, both simulations qualitatively captured most of the up/down regulation trends. Fig. S4 shows the metabolic network (modified from the Escher [28] map for the *E. coli* core model) with reactions of Fig. 2(b) highlighted and color-coded according to the gene expression data. Finally, we reproduced the other experimental scenarios from Enjalbert *et al.* [22] with the uniform population model, adjusting only the initial values of biomass, glucose and acetate. We observed that when high acetate concentrations are present in the medium a uniform shift can well reproduce the biomass profile (Fig. S5(c)), while this is not the case when only low acetate concentrations are available (Fig. S5(b)).

**Figure 1:**
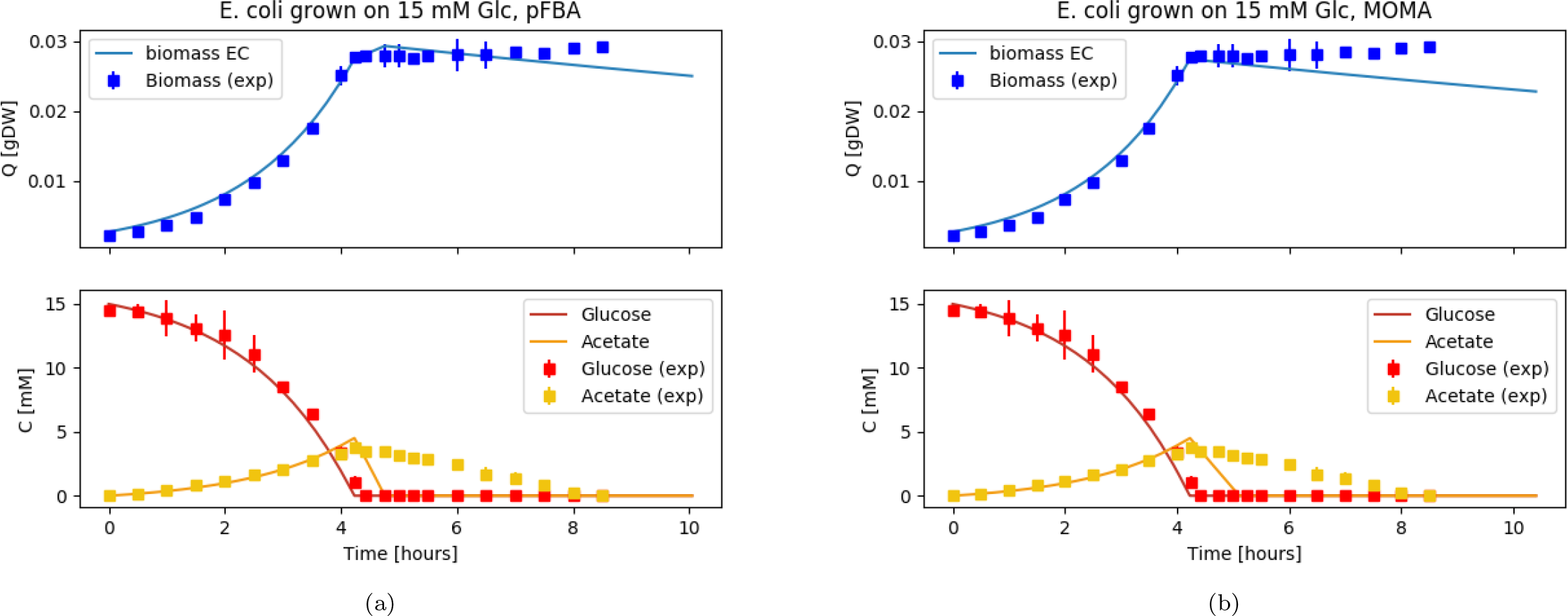
Diauxic growth of *E. coli* modeled as a uniform population in batch conditions. Simulation (lines) is compared to data (squares) from Enjalbert *et al.* [22] as a function of time. Biomass (blue, top sub-plots), glucose and acetate (red and yellow, bottom sub-plots) are shown. The flux distribution at each time step is obtained with pFBA (a) or MOMA (b).

**Figure 2:**
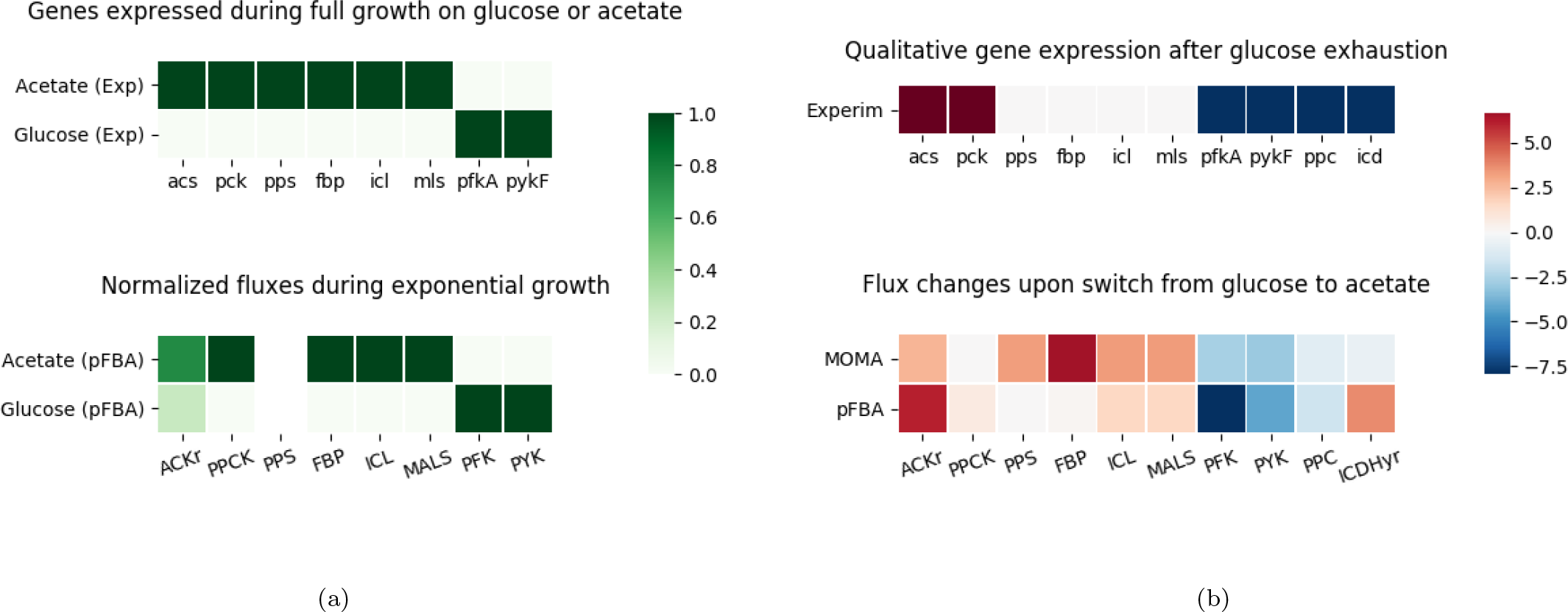
Comparison of experimental information on gene expression levels with simulated flux distributions. The top plots represent qualitatively the gene expression data from Enjalbert *et al.* [22]. Flux solutions in the simulations for the reactions associated to the reported key genes are compared (a) between two independent simulations with *E. coli* exponentially growing either on acetate or on glucose; (b) within the same simulation (Fig. 1, growth on glucose simulated with MOMA or pFBA) before and after the point of glucose exhaustion.

**Table 1:** Fixed parameters for all simulations. The lower bound for oxygen exchange as well as the upper bound for acetate exchange are calibrated on the data from Varma and Palsson [18]. The death rate *δ* is computed assuming a cell death of 1% per generation [44] and a generation time of 20 min. The Michaelis-Menten parameters for substrate uptake are taken from Gosset [45]. These parameters were also used in previously published dFBA implementations [46].

| | **L.B.**<br>**EX_O2**<br>$\frac{\text{mmol}}{\text{gDW hr}}$ | **U.B.**<br>**EX_Ac**<br>$\frac{\text{mmol}}{\text{gDW hr}}$ | $\delta$<br>$\text{hr}^{-1}$ | $V_M^{\text{Glc}}$<br>$\frac{\text{mmol}}{\text{gDW hr}}$ | $K_M^{\text{Glc}}$<br>mM | $V_M^{\text{Ac}}$<br>$\frac{\text{mmol}}{\text{gDW hr}}$ | $K_M^{\text{Ac}}$<br>mM |
| --- | --- | --- | --- | --- | --- | --- | --- |
| Value | -11.5 | 3.0 | 0.03 | 10.0 | 0.01 | 10.0 | 0.01 |

**Table 2:** Parameters of the simulations.

| | **BM(0)**<br>$10^{-3}$ gDW | $\left. \frac{\text{EC}_{\text{Glc}}}{\text{EC}_{\text{Ac}}} \right\vert _0$ | **Glc(0)**<br>mmol | **Ac(0)**<br>mmol | $\xi$<br>$\frac{\text{mmol}}{\text{hr}}$ | $\zeta$<br>$\frac{\text{mmol}}{\text{hr}}$ | $t_x$<br>hr | $\psi_0$<br>$\text{hr}^{-1}$ | $\mathbf{V}_M^\psi$<br>$\text{hr}^{-1}$ | $\mathbf{K}_M^\psi$<br>mM | $\phi_0$<br>$\text{hr}^{-1}$ | $\mathbf{V}_M^\phi$<br>$\text{hr}^{-1}$ | $\mathbf{K}_M^\phi$<br>mM | $\varepsilon$ |
| --- | --- | --- | --- | --- | --- | --- | --- | --- | --- | --- | --- | --- | --- | --- |
| Fig. S1(a), S1(c) | 0.3 | NA | 10.8 | 0.4 | 0.0 | 0.0 | 0.0 | 0.0 | 0.0 | 0.0 | 0.0 | 0.0 | 0.0 | 0.0 |
| Fig. S1(b), S1(d) | 0.24 | NA | 0.82 | 0.1 | 1.1 | 0.0 | 0.0 | 0.0 | 0.0 | 0.0 | 0.0 | 0.0 | 0.0 | 0.0 |
| Fig. 1(a), 1(b), S5(a) | 2.7 | NA | 15.0 | 0.0 | 0.0 | 0.0 | 0.0 | 0.0 | 0.0 | 0.0 | 0.0 | 0.0 | 0.0 | 0.0 |
| Fig. 3(a) | 2.7 | $\frac{0.95}{0.05}$ | 15.0 | 0.0 | 0.0 | 0.0 | 0.0 | 0.0 | 0.0 | 0.0 | 0.0 | 0.0 | 0.0 | 0.0 |
| Fig. 3(b), S5(d) | 2.7 | $\frac{0.95}{0.05}$ | 15.0 | 0.0 | 0.0 | 0.0 | 0.0 | 0.04 | 0.0 | 0.0 | 0.04 | 0.0 | 0.0 | 0.9 |
| Fig. S5(g) | 2.7 | $\frac{0.95}{0.05}$ | 15.0 | 0.0 | 0.0 | 0.0 | 0.0 | 0.04 | 0.2 | 30.0 | 0.04 | 0.2 | 5.0 | 0.9 |
| Fig. S5(b) | 3.8 | NA | 15.0 | 0.0 | 0.0 | 9.1 | 4.0 | 0.0 | 0.0 | 0.0 | 0.0 | 0.0 | 0.0 | 0.0 |
| Fig. S5(e) | 3.8 | $\frac{0.75}{0.25}$ | 15.0 | 0.0 | 0.0 | 9.1 | 4.0 | 0.04 | 0.0 | 0.0 | 0.04 | 0.0 | 0.0 | 0.9 |
| Fig. 4(a), S5(h) | 3.8 | $\frac{0.75}{0.25}$ | 15.0 | 0.0 | 0.0 | 9.1 | 4.0 | 0.04 | 0.2 | 30.0 | 0.04 | 0.2 | 5.0 | 0.9 |
| Fig. S5(c) | 6.0 | NA | 15.0 | 32.0 | 0.0 | 9.1 | 4.0 | 0.0 | 0.0 | 0.0 | 0.0 | 0.0 | 0.0 | 0.0 |
| Fig. S5(f) | 6.0 | $\frac{0.75}{0.25}$ | 15.0 | 32.0 | 0.0 | 9.1 | 4.0 | 0.04 | 0.0 | 0.0 | 0.04 | 0.0 | 0.0 | 0.9 |
| Fig. 4(b), S5(i) | 6.0 | $\frac{0.75}{0.25}$ | 15.0 | 32.0 | 0.0 | 9.1 | 4.0 | 0.04 | 0.2 | 30.0 | 0.04 | 0.2 | 5.0 | 0.9 |
| M9G (m.c.) | 2.7 | $\frac{0.95}{0.05}$ | 15.0 | 0.0 | 0.0 | 0.0 | 0.0 | 0.04 | 0.2 | 30.0 | 0.04 | 0.2 | 5.0 | 0.9 |
| M9GA (m.c.) | 6.0 | $\frac{0.75}{0.25}$ | 15.0 | 32.0 | 0.0 | 0.0 | 0.0 | 0.04 | 0.2 | 30.0 | 0.04 | 0.2 | 5.0 | 0.9 |

### *E. coli* diauxie modeled with a mixed population

We used two GEMs (same parameter values as before) to model *E. coli* monoculture as a mixture of two populations, one adapted to grow on glucose and one adapted to grow on acetate. The two models EC_Glc_ and EC_Ac_ are hence constrained to exclusively take up the corresponding carbon source. Two transition functions, dependent on acetate or glucose concentrations, are introduced to model cellular differentiation and shift from one population to the other (see Materials and Methods for details). We ran simulations to compare the different scenarios investigated experimentally by Enjalbert *et al.* [22]. Initial values of biomass, glucose and acetate were adjusted to the corresponding datasets. The transition rates, as well as the initial population ratios, were chosen following the assumption, supported by a simple mathematical model, that in constant environments the populations will converge to a constant ratio (see Text S1 for details). Fig. 3 shows simulations for the same condition as Fig. 1(a), with same absolute initial biomass, distributed in this case as 95% EC_Glc_ and 5% EC_Ac_. This initial ratio was chosen considering the range of steady-state values for the population ratio (reported in Table S1) as well as considering that it is reasonable to assume that a higher number of cells will be adapted to grow on glucose, which is the carbon source on which laboratory cultures are usually maintained. Fig. 3(a) shows the simulation results for a scenario without transitions between the two states, whereas the results of Fig. 3(b) were obtained with active transition functions, defined here by constant transition rates as reported in Tab. 2. Although both Fig. 3(a) and Fig. 3(b) well capture the biomass (*R*^2^ = 0.987 and *R*^2^ = 0.990, respectively) and glucose concentrations (*R*^2^ = 0.996 and *R*^2^ = 0.997, respectively), only the simulation including the population transition realistically reproduced the acetate consumption (*R*^2^ = 0.336 and *R*^2^ = 0.951 respectively), as well as a lag phase before culture crash. Neither of the two simulations captured the eventual recovery of growth hinted by the last data points. We reproduced two other results (where only biomass measurements are available) from Enjalbert *et al.* [22], again using the same GEMs and changing only the initial conditions (biomass quantity and distribution among EC_Glc_ and EC_Ac_) and the experimental setup accordingly. By modeling the population transition with the same constant rate, we were able to explain the biomass profile in the case where *E. coli* is grown on 15 mM glucose and after glucose exhaustion the acetate concentration is maintained at around 4 mM (Fig. S5(e), *R*^2^ = 0.986), but not in the case where *E. coli* is grown on 15 mM glucose and 32 mM acetate, and after glucose exhaustion the acetate concentration is maintained at the same high level (Fig. S5(f), *R*^2^ = 0.727). We therefore introduced a dependency of the transition functions on the substrate concentration (see Materials and Methods for details) that well captures all the experimental scenarios with a unique set of parameters (Fig. S5(g), S5(h), S5(i)). Fig. 4(a) shows that an *E. coli* population starting with 95% EC_Glc_ and 5% EC_Ac_ well describes the biomass dynamics (*R*^2^ = 0.985) and the glucose exhaustion point after around 4 h when acetate is maintained at 4 mM. Again, without fine-tuning the GEM simulation parameters, Fig. 4(b) shows that an *E. coli* population starting with 75% EC_Glc_ and 25% EC_Ac_ reproduces the biomass measurements (*R*^2^ = 0.940) and the glucose exhaustion point after around 4 h also in the experimental setup with acetate maintained at 32 mM. The effect of varying the initial biomass ratios in the different experimental conditions is shown in Fig. S6. Overall, simulations starting with 95% EC_Glc_ and 5% EC_Ac_ or 75% EC_Glc_ and 25% EC_Ac_ did not show strong differences, but further reducing the percentage of EC_Glc_ (and leaving the range of steady-state values of Table S1) resulted in drastic changes to the shape of the growth curves. The initial condition of 75% EC_Glc_ and 25% EC_Ac_ population distribution for Fig. 4(b) is also justified by a difference in the experimental initial values for the biomass quantity (see Fig. S7).

**Figure 3:**
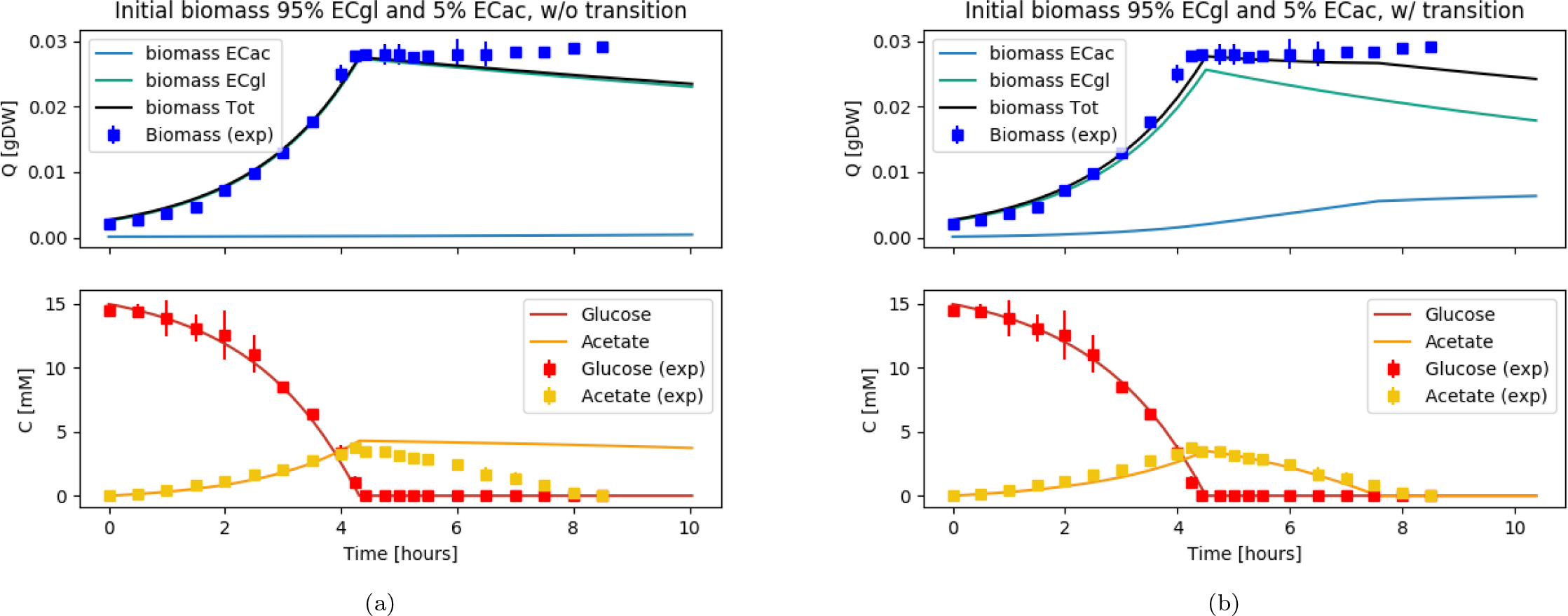
Diauxic growth of *E. coli* modeled as a mixture of two *E. coli* populations EC_Glc_ and EC_Ac_ growing exclusively on glucose or acetate, respectively, and without (a) or with (b) possibility to shift from one population to the other. Simulation (lines) is compared to data (squares) from Enjalbert *et al.* [22] as a function of time. The upper plots show simulation results (using pFBA) for EC_Glc_ and EC_Ac_ biomasses (light blue and aqua, respectively) and the observable *E. coli* biomass (black line simulation, blue dots data). The bottom plots show glucose and acetate (red and yellow respectively).

**Figure 4:**
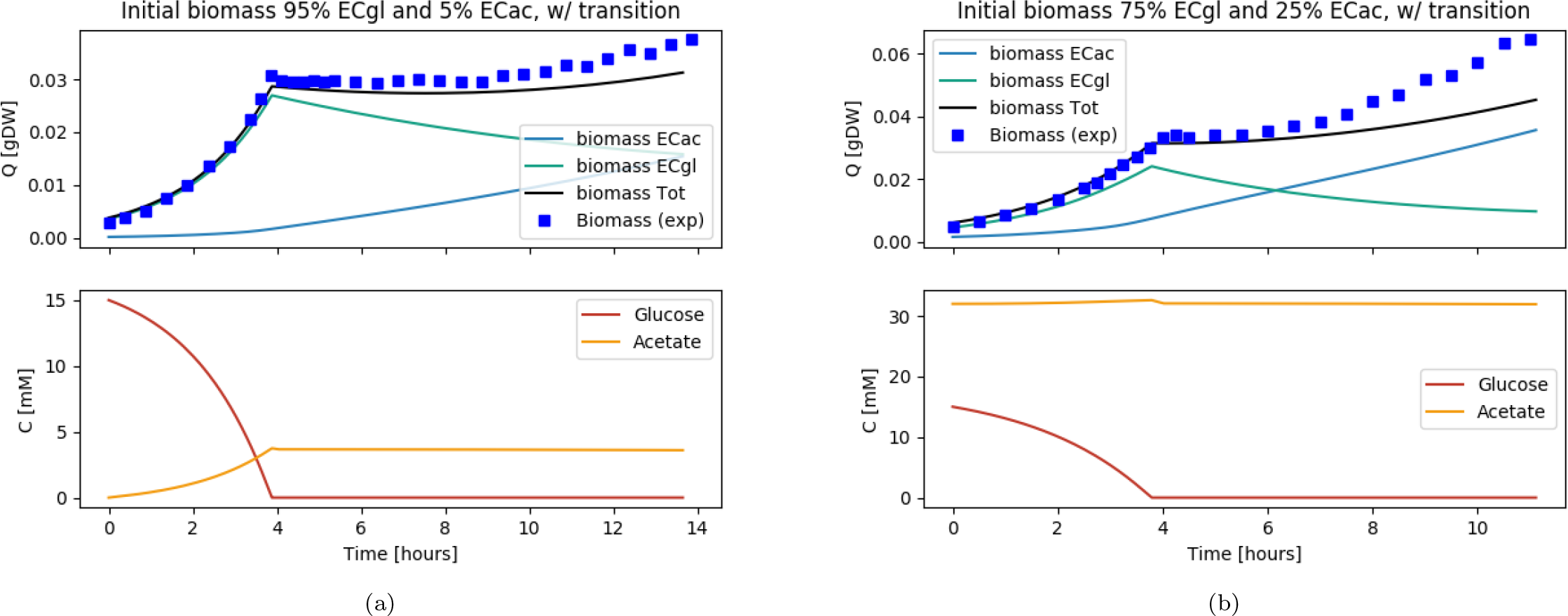
Diauxic growth of *E. coli* modeled as a mixture of two *E. coli* populations EC_Glc_ and EC_Ac_ growing exclusively on glucose or acetate, respectively, with possibility to shift from one population to the other. (a) *E. coli* grows on 15 mM glucose and after glucose is exhausted, acetate concentration is kept at about 4 mM. (b) *E. coli* grows on 15 mM glucose and 32 mM acetate and after glucose is exhausted, acetate concentration is maintained at around that concentration. The upper plots show simulation results (using pFBA) for EC_Glc_ and EC_Ac_ biomasses (light blue and aqua lines, respectively) and the observable *E. coli* biomass (black line simulation, blue dots data from Enjalbert *et al.* [22]). The bottom plots show simulation results for glucose and acetate (red and yellow lines respectively).

### Lag time for growth on acetate explained with population distribution

Enjalbert *et al.* [22] showed different trends in the lag time of *E. coli* cultures required to achieve maximal growth after GE. In their switch experiments, they sampled at different time points “mother cultures” of *E. coli* cells growing in batch conditions on 15 mM glucose alone (M9G condition) or on 15 mM glucose and 32mM acetate (M9GA condition), and re-inoculated the sampled cells as “daughter cultures” into fresh medium exclusively containing glucose (M9G condition) or acetate (M9A condition). We replicated this experiment *in silico* by running first simulations under the M9G and M9GA conditions. For the M9G mother culture, we used the simulation of the mixed EC_Glc_ and EC_Ac_ population shown in Fig. 3(b), because the experimental conditions are the same. For the M9GA mother culture, we did not have an experimental reference dataset and we simulated a new scenario similar to that shown in Fig. 4(b), with the same initial population composed of 75% EC_Glc_ and 25% EC_Ac_, but without the feeding of additional acetate. The GE time point is about 4.6 h for M9G and 3.9 h for M9GA, consistent with the observations of Enjalbert *et al.* [22] (data not shown). The *in silico* mother cultures were sampled at regular time intervals to obtain the initial biomass distribution of EC_Glc_ and EC_Ac_ for the daughter cultures (reported in Tab. 3) and the lag time of each daughter culture was computed (see Materials and Methods for details). Fig. 5 shows the simulation results compared with the experimental data from Enjalbert *et al.* [22]. The error bars on the simulated lag time were obtained by varying the initial biomass ratio of the daughter cultures by ±15%. A quantitative agreement between simulation and experimental results was only achieved in the M9G-M9G switch experiment (Fig. 5(a)) with the correct prediction of almost zero lag time for the daughter cells, but the trend for the delay to reach maximal growth was in general qualitatively reproduced also for the other scenarios. According to the simulations, cultures switched from M9G to M9A (Fig. 5(a)) need about 1.5 h before reaching maximal growth, which is more than twice the duration observed experimentally. For cultures pre-grown in M9GA (Fig. 5(b)), we observed both in simulations and in experiments a decreasing lag time for daughter cultures sampled after GE for the M9GA-M9A switch, and an increasing lag time for the M9GA-M9G switch. Additional studies are reported in Fig. S8. In particular, Fig. S8(a-d) shows the dependence of the lag time in the daughter cultures on the maximal transition values and Fig. S8(e-h) shows the same dependence, including also the distribution of the biomass ratio in the mother cultures, for a limited set of parameters.

**Figure 5:**
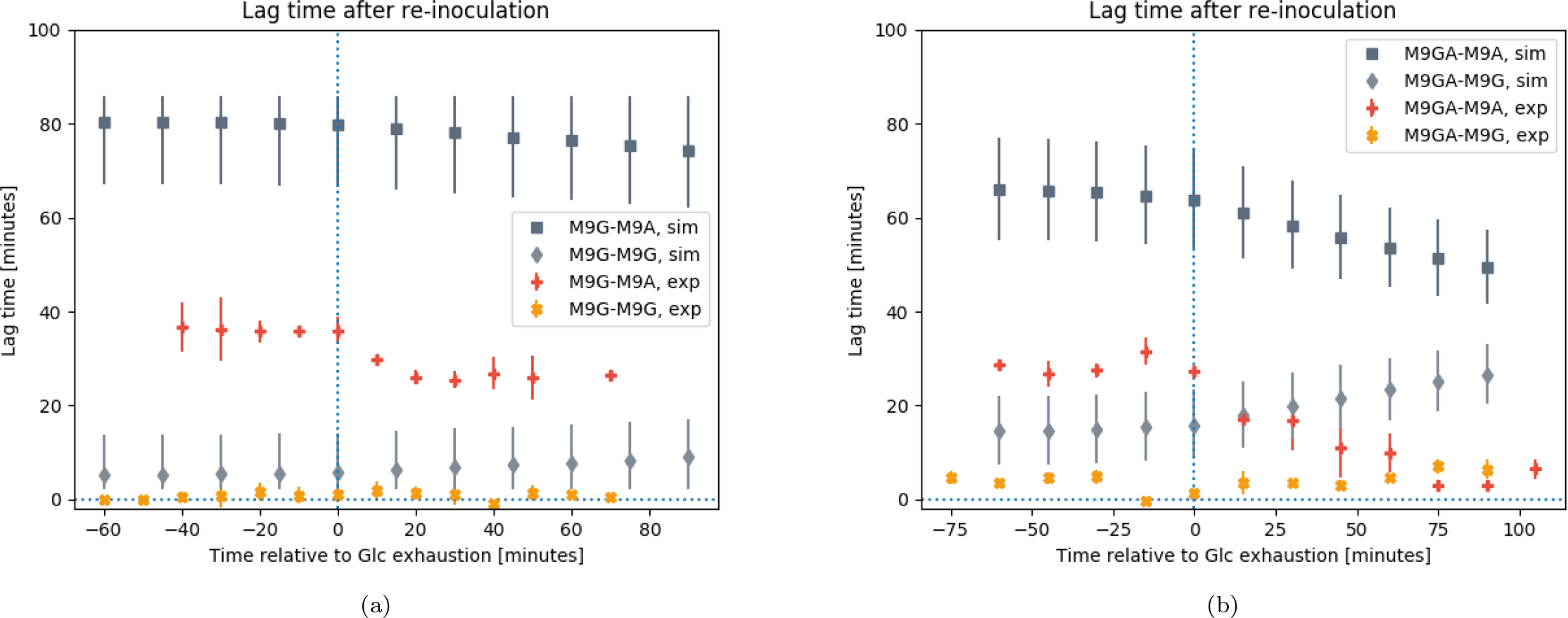
Simulation (dark and light gray points) and experimental (orange and yellow points, data from Enjalbert *et al.* [22]) results for the delay of daughter cultures before reaching maximal growth after media switch. Mother cultures are grown either on 15 mM of glucose (M9G, (a)) or 15 mM of glucose and 32 mM of acetate (M9GA, (b)). Daughter cultures are re-inoculated into fresh media with either 15 mM of glucose (M9G, square and plus markers) or 45 mM of acetate (M9A, diamond and cross markers). The simulation error bars are obtained by varying the initial population ratios (obtained from sampling the simulated mother cultures) by ±15%.

**Table 3:** Percentage of EC_Glc_ biomass in M9G and M9GA conditions at time points relative to glucose exhaustion, in hours.

|  | -1.0 | -0.75 | -0.5 | -0.25 | 0 | 0.25 | 0.5 | 0.75 | 1.0 | 1.25 | 1.5 |
| --- | --- | --- | --- | --- | --- | --- | --- | --- | --- | --- | --- |
| M9G | 95.1 | 95.0 | 94.8 | 94.5 | 94.3 | 93.2 | 92.3 | 91.4 | 90.6 | 89.4 | 88.2 |
| M9GA | 78.7 | 78.6 | 78.1 | 77.3 | 76.5 | 73.2 | 70.1 | 67.2 | 64.5 | 62.0 | 59.7 |

## 3 Discussion

We have investigated a fundamental example of metabolic adaptation, namely the diauxic growth of *E. coli* on glucose and acetate, aiming to test whether a dynamic metabolic modeling approach can capture diauxie in monocultures of *E. coli* as the observable emergent result of individual (sub-population) behavior. To this end, we first developed a modeling framework to integrate dynamic models (ODE systems) with structural models (metabolic networks) and then performed simulations to reproduce published experimental results *in silico*.

### Avoiding fine-tuning of model parameters

One recurrent criticism of stoichiometric and constraint-based modeling approaches, such as FBA, is that they can easily be adjusted to reproduce experimental results by *ad hoc* changes of flux constraints. Indeed, we observed that a condition-specific fine-tuning of the constraint on acetate uptake could reproduce fairly well the growth dynamics of the different experiments (data not shown). However, the change of such constraint from one experimental condition to another is not biologically justified. Although some extensions of the FBA approach such as FBA with molecular crowding (FBAwMC [29]) provide reasonable ways to constrain the metabolic fluxes and were shown to reproduce carbon consumption hierarchies, they also require extensive parameterization. We therefore chose to use the basic FBA approach, limiting the number of constraint imposed and with parameters mostly from experimental measurements (Tab. 1). In the case of oxygen uptake and acetate secretion, we calibrated the constraints using the data from Varma and Palsson [18], where an *E. coli* diauxic shift from glucose to acetate was first simulated using a genome scale model. The FBA parameters were left unchanged to reproduce the independent experiments reported by Enjalbert *et al.* [22]. The use of an independent set of data to calibrate the FBA model parameters is a possible way to improve the confidence in subsequent results. Further model parameters of the ODE system were chosen according to reasonable hypotheses, and were adjusted slightly to achieve fair agreement with the experimental results, consistently among all the simulations. The initial conditions were specific to the experiments we aimed to reproduce.

### Standard dFBA allows for abrupt metabolic readjustments

The flux distributions obtained from FBA solutions represent an average picture of the metabolic state of a population, which is in general modeled using a single genome scale model. Therefore, standard dFBA implementations, in which the FBA constraints evolve according to the updated external conditions, will reproduce the average change in metabolic state of the population in response to external variations. This is equivalent to assuming that a population undergoes a coordinated, uniform metabolic shift under changing environmental conditions. Furthermore, such transitions are generally abrupt with dFBA models. We therefore tested two alternative approaches to simulate the diauxic shift in uniform *E. coli* monocultures, solving the FBA problems either with pFBA (mostly equivalent to the usual dFBA implementations) or with an adaptation of the MOMA algorithm. In the latter case, instead of minimizing the difference in flux distribution between a “wild-type” GEM and a modified one (original MOMA implementation), we used the same concept to integrate the dFBA system while also imposing the following condition: at time *t*_*i*_ the flux solution differs minimally from that at time *t*_*i*−1_, where the time steps are set by the integration routine. Contrary to our expectations however, this approach did not achieve smoother metabolic adjustments in the system in response to the changing external conditions. Instead, both implementations resulted in abrupt changes in the flux distributions following the shift from glucose to acetate metabolism (Fig. S3). More sophisticated implementations of a dynamic MOMA model (*e.g.* computing the minimal adjustment based on a subset of biologically relevant variables) might succeed in achieving smooth metabolic transitions, but will require the introduction of additional parameters and *ad hoc* biological hypotheses. In a similar way, biologically justified extensions of FBA such as FBAwMC [29] might provide better descriptions of an average and uniform population-level metabolism, but typically need the empirical determination of large numbers of organism-specific parameters.

### Monocultures can be modeled as multi-sub-population systems to capture individual heterogeneity

With the introduction of two basic assumptions (first, there are two distinct metabolic states consuming either glucose or acetate; second, transition from one state to the other is driven by glucose and acetate concentrations) we were able to capture all the experimental trends published by Enjalbert *et al.* [22] with the same computational model. The transitions between two states were modeled as Hill functions of the corresponding substrate concentrations with a noise offset representing a constant, small noise component in cell regulation. Although other transition laws could have been chosen, Hill functions conveniently model concentration-dependent shifts between two states. For example, if acetate is highly abundant, more cells in the glucose-consumption state will shift to the acetate-consumption state in response to the change in environment. Finally, the introduction of a transition efficiency term was motivated by the observation that cells can get “lost in transition”, an effect that was estimated to account for the death of ~7% of yeast cells, which cannot initiate glycolysis following a shift to high glucose levels [30]. Using a simple mathematical model (Text S1) we identified ranges for the parameters of the transition functions and selected reasonable values that would return good agreement between simulations and experiments. Both values for the constant transition rate (4% h^−1^) and for the maximal transition rate (20% h^−1^) were in good agreement with measured average protein turnover rates in *E. coli* cultures from the literature [31, 32, 33]. Simulation results were mostly in very good agreement with the experimental data and our results strongly further support the idea, suggested over the last few years by independent research on different organisms [21, 25, 34], that monocultures are an ensemble of sub-populations in different metabolic states, partially regulated by the environmental conditions. When the acetate concentrations were too low to support growth, it was sufficient to model the transition as a constant random process. In contrast, in order to reproduce the data under conditions with high acetate concentrations, we needed to introduce an active transition rate dependent on substrate concentrations. Interestingly, this assumption alone is sufficient to model the experimentally observed growth rate, without further fine tuning of model parameters. The introduction of substrate-dependent transition functions is also consistent with the experimental observations of Kotte *et al.* [27], supporting the hypothesis that a monoculture undergoes diversification in response to environmental changes.

### The lag phase between growth on different substrates can be explained by population distributions

With standard dFBA simulations, the metabolic transition during the shift from one carbon source to another is abrupt, and no lag phase is observable. This is rarely the case and, most remarkably, the duration of the lag phase between the exhaustion of the favored carbon source and the resumption of optimal growth on the alternate carbon source is highly variable under different environmental conditions. This observation can easily be explained as an emergent property of sub-population dynamics. Our simulations are consistent with the explanation that the delay in the resumption of full growth actually depends on the relative abundance of the two sub-populations. Although the simulation results did not reproduce the experimental data quantitatively, all qualitative trends were fully explained. Several factors may explain these discrepancies. For example, the lack of experimental data concerning the mother cultures (in terms of biomass, glucose and acetate dynamics) made it impossible to calibrate the initial model population. This could introduce a significant bias in the later sampling and determination of the sub-population ratio, thus strongly influencing the quantification of the lag time, which is highly correlated with the population distribution (Fig. S8). Solopova *et al.* [25] showed that the density of a *Lactococcus lactis* population (translating in practice to the rate at which the primary carbon source was consumed) played a significant role in determining the proportion of cells successfully transitioning to growth on the secondary carbon source. The connection between lag time and subpopulation distribution could in principle be exploited to estimate initial population distributions from lag time measurements. However, with the currently available data it is difficult to assess the robustness and reliability of such predictions, and further investigation is therefore required, including devoted experiments to determine initial conditions. An additional source of discrepancy between our quantitative results and the experimental measurements could be the experimental procedure itself. For example, abrupt changes in conditions, such as the re-inoculation of daughter cultures into a different medium in the switch experiments might select for additional adaptation strategies. Interestingly, we observed a dramatic improvement in the quantitative agreement between experiment and simulation by relaxing the condition imposing no growth for populations inoculated on the “wrong” carbon source (data not shown). By allowing the glucose-consuming population sampled from glucose mother cultures to growth more slowly on acetate, we mimick a situation in which cells store resources and are able to survive a bit longer. On the other hand, allowing reduced growth on acetate (glucose) for the glucose-consumer (acetate-consumer) population that was exposed to both carbon sources in the mixed mother cultures could be a proxy for a memory effect. Bacterial cells do show memory effects upon changes in environmental conditions [26], but to explore this potential explanation further, more systematic experiments would be necessary to carefully and reproducibly determine the lag times as functions of external parameters. Finally, a recent stochastic model of the regulatory network of diauxic growth in *E. coli* suggests that the limitations of biological sensors are responsible for the lag phase [35]. From these results we can infer that in our model the transition functions, currently depending on the absolute concentration of one carbon source at a time, might not be able to capture the fine details of population shifts. A possible extension would be to introduce more complex transition mechanisms dependent on relative concentrations of primary and secondary carbon sources, a process that would need dedicated experiments for the construction and validation of the new transition functions.

### Sub-populations in the dynamic metabolic modeling approach

We developed a modeling framework to perform FBA simulations embedded in a system of ODEs. Building on previous methods and approaches [19, 36], we further extended the standard dFBA implementation and introduced novel concepts. In particular, standard dFBA approaches assume that fluxes can instantaneously change to adapt to new environmental conditions, and flux solutions at subsequent time steps might differ significantly. This is an obvious limitation when aiming to capture diauxic shift, where lag phases, highly dependent on the environmental conditions, are typically observed. We implemented the MOMA algorithm (originally developed to model the response to genetic perturbations in static FBA) in dFBA to minimize the metabolic adjustment between different time points. Furthermore, we integrated dynamic mechanisms into dFBA that cannot be included in metabolic models, such as population transitions. Indeed, although the use of dFBA to model sub-populations bears some similarities to other platforms for the simulation of microbial communities, a notable difference in our formulation is the capacity of sub-populations to interconvert. The current study relied on the *a priori* knowledge that only two carbon sources would be available to *E. coli*, thus motivating the development of a two sub-population community, but in principle an arbitrary number of sub-populations can be defined, and more generic transition functions introduced. Further experiments, in particular single-cell studies, could be designed to define and parameterize these transition functions. Thanks to the object oriented design of the framework, it is relatively easy to introduce other functions regulating the constraints on specific reaction fluxes in the FBA problem. In this way, different hypotheses can be extensively tested to better understand how to capture regulatory dynamics in dFBA. Notably, the methods developed in this framework to study population heterogeneity could then be transferred to other platforms more specific for microbial community modeling where different features are implemented (*e.g.* spatial structure [19] or community-level objectives [37]). Finally, the framework could also be developed further to include stochastic mechanisms, such as mutations that would alter the function of metabolic genes. Indeed, our implementation of the dFBA algorithm is able to call different methods at each time step, *e.g.* to update the flux rates, and a regulatory function with random components could be in principle defined.

### Outlook

There is extensive experimental evidence that bacteria differentiate into sub-populations as a result of survival strategies [25, 27]. Simulations based on standard dFBA model the dynamics of cells by predicting the putative average behavior of a whole population. For example, if a population of cells globally utilizes a combination of two carbon sources, dFBA would predict metabolic states in which both carbon sources are utilized simutaneously. Our model assumes that cells are either in the glucose or acetate consuming state, with an instantaneous transition between these two sub-populations that follows a simplistic rule which cannot capture intermediate states. This simplification is both practical and plausible when we observe population dynamics as the emergent properties of individual behavior, and it works well in dynamically changing environments with a continuous transition. However, rather than having a well-defined metabolic state, especially during the transition between states, cells might exhibit a mixed state, which could be described as a superposition of ‘pure’ states, analogous to the state vectors in quantum physics. Furthermore, our approach suggests a fundamental difference in the strategies to account for metabolic fluxes in heterogeneous populations, because the average fluxes in a uniform population might differ from the cumulative average fluxes of sub-populations. Further investigations of this novel concept of superimposed metabolic states will provide a promising new approach to study the principles of metabolic regulation.

## 4 Materials and Methods

### FBA methods

In stoichiometric models, the stoichiometric matrix *S*(*m* × *n*) is defined with the row and column dimensions corresponding to the numbers of metabolites *m* and reactions *n* respectively, the elements *s*_*ij*_ being the stoichiometric coefficients of metabolite *i* taking part in reaction *j*. FBA defines and solves the following LP problem:

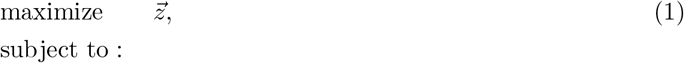

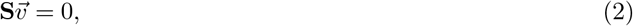

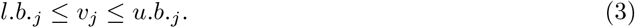

The steady-state assumption (Eq. 2) gives a system of equations that is under-determined and has an infinite number of solutions. Constraints on the fluxes (Eq. 3) allow us to restrict the solutions to a convex solution space, but still result in an infinite number of solutions. The definition of an objective (Eq. 1) selects one solution, but generally this is still not unique for large (genome-scale) metabolic networks.

We consider two extensions to the FBA problem definition, namely pFBA [14] and MOMA [15]. We then use these two methods to solve the FBA problem in an approach similar to dFBA [16]. Assuming that metabolism evolves towards the efficient utilization of resources, pFBA finds the minimal flux distribution that returns the same objective defined by the FBA problem. We use the pFBA implementation from COBRApy [38] with maximal flux through the biomass reaction as the objective function. Considering that metabolism must respond quickly to perturbations, MOMA implements a quadratic algorithm to find the FBA solution after gene deletion that is most similar to the optimal wild-type configuration. In our case, we do not introduce modifications to the metabolic network but rather require that the MOMA solution obtained at time *t*_*i*−1_ is used to compute the MOMA solution at time *t*_*i*_ as the minimally different solution that satisfies the objective function. Also here the objective function is maximal flux through the biomass reaction. We use the MOMA implementation from COBRApy [38] in the linear approximation, with a slight modification to allow the LP problem to be reset in an iterative manner, which is necessary to run MOMA within the dFBA approach.

### Modeling framework integrating ODE and FBA

In the SOA of dFBA the boundary conditions in Eq. 3 are updated at discrete time steps according to the solution of the FBA problem in the previous time interval. Assuming quasi-steady-state conditions, *i.e.* that metabolism readjustments are faster than external environmental changes, dFBA can approximate the dynamic response of a GEM to a changing environment. Our approach is an extension of dFBA. The model is built as a system of ODEs, whose dimension depends on the dynamics to be modeled. Each ODE describes the variation in time of biomass, metabolites, or other regulatory/dynamic processes. The biomasses and the metabolites can, but do not necessarily, be linked to the corresponding variables in a GEM. Their ODEs vary according to a function that can then depend on the flux solutions 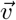

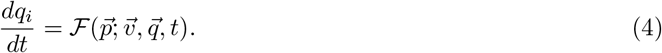

The ODE system is then solved using integration routines with an automated choice of time step. Each integration step solves the FBA problem (or pFBA, or MOMA) to obtain the current reaction rates for Eq 4, updates the metabolite quantities according to the FBA solution, re-computes the flux boundaries of Eq. 3 according to specific reaction kinetics (typically Michaelis-Menten enzyme kinetics), and redefines the FBA problems with the new boundaries and/or other regulatory mechanisms defined by the user.

The modeling framework is written in Python (Python Software Foundation, https://www.python.org/) following the object-oriented programming (OOP) paradigm for efficiency and flexibility. The framework uses functionality from the following third-party packages: numpy [39], scipy [40], matplotlib [41], CO-BRApy [38], and pandas [42]. In particular, we use COBRApy methods to solve the FBA problems and Python integrators from the scipy.integrate method ode to solve the system of ODEs.

### *E. coli* uniform population model

We used a previously reported core version of *E. coli* GEM [43] downloaded from http://bigg.ucsd.edu/models/e_coli_core. The *E. coli* model EC_any_ is constrained on the consumption of “any” carbon source (glucose, Gl, or acetate, Ac) solely by the environmental conditions, and the lower bound of the exchange reactions (EX_Glc_e and EX_Ac_e respectively) follows two simple Michaelis-Menten kinetics:

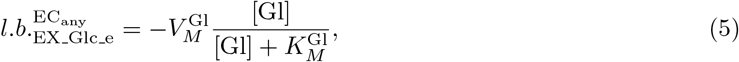

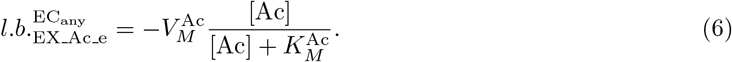

The ODE system is defined as

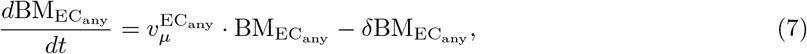

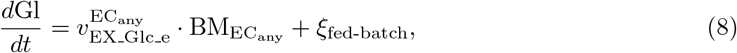

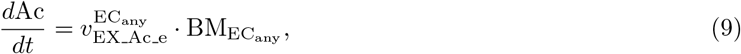

where *v*_*μ*_ is the reaction rate of the biomass function (proxy for growth rate) in the FBA model, *δ* is the cell death rate and ξ_fed-batch_ is a positive rate under fed-batch conditions and zero under batch conditions. Parameters and initial conditions are summarized in Tab. 2. Either pFBA or MOMA can be used to solve the FBA problem.

### *E. coli* mixed population model

Two *E. coli* core models are loaded and defined as either Glucose consumer (EC_Glc_) or Acetate consumer (EC_Ac_) by switching off uptake of the other carbon source

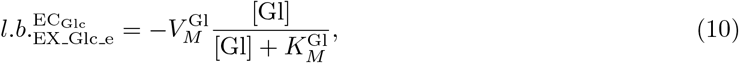

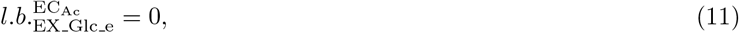

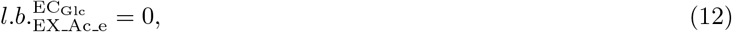

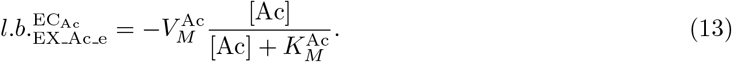

The ODE system is defined as

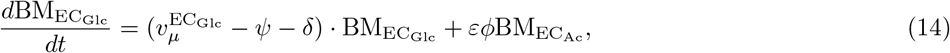

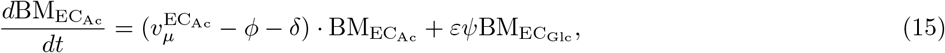

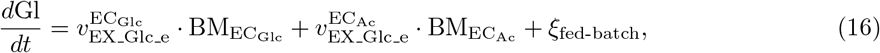

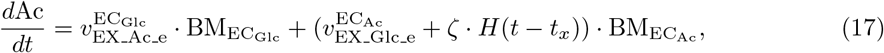

where ζ · *H*(*t* − *t*_*x*_) is a Heaviside function activated at the time *t*_*x*_ of glucose exhaustion in order to keep acetate constant, and *ψ* and *ϕ* are functions that model the cellular shift from EC_Glc_ to EC_Ac_ and EC_Ac_ to EC_Glc_, respectively, and 0 < *ε* < 1 is a positive factor representing the transition efficiency. The functions *ψ* and *ϕ* are modeled as Hill functions with a noise offset

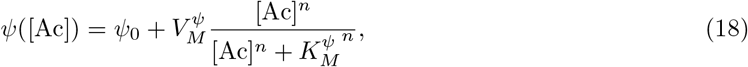

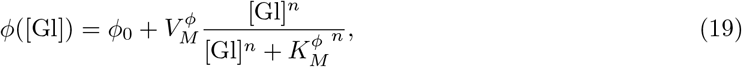

and for 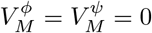 they are constant transition rates. For the simulations presented herein, we used a Hill coefficient *n* = 5. Indeed, simulations seem to work best for a transition function with a high degree of cooperativity, and the results are robust with respect to small deviations relative to this value. The other parameters and initial conditions, specific to the different simulations, are summarized in Tab. 2. For mixed-population simulations, pFBA is used to solve the FBA problem.

### Switch experiment simulations

Two *E. coli* mixed population model simulations are run as “mother cultures” as shown in Tab. 2 for “M9G” and “M9GA” conditions (glucose and glucose plus acetate, respectively). From each mother culture we sample 11 time points between −1 and +1.5 h from the corresponding GE time (4.6 h for M9G and 3.9 h for M9GA) to obtain the biomass ratio between EC_Glc_ and EC_Ac_ used as the initial condition for the re-inoculation simulations. The percentage of EC_Glc_ biomass at these time points is shown in Tab. 3. The “daughter cultures” are then grown under “M9G” glucose-only or “M9A” acetate-only conditions (see Tab. 2), yielding 44 simplified simulations, 11 for each of the following 4 switch experiments: M9G to M9G; M9G to M9A; M9GA to M9G; M9GA to M9A. For each simulation, the lag time is computed according to Enjalbert *et al.* [22]:

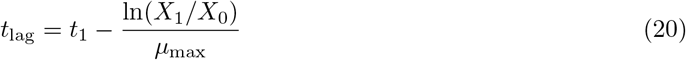

where *X*_0_ is the total initial *E. coli* biomass, *X*_1_ is the total *E. coli* biomass value at time *t*_1_ (1.5 h as in [22]), and *μ*_max_ values are used according to Enjalbert *et al.* [22].

## Published experimental data

Experimental data (values with standard deviations, when available) from Enjalbert *et al.* [22] were kindly provided by Prof. Enjalbert. The data from Varma and Palsson [18] were extracted from the original publication using WebPlotDigitizer [47].

## Availability of data and materials

The version of the modeling framework used to obtain the results presented in this manuscript (v1.1) is publicly available with instructions to install and run simulations at https://github.com/QTB-HHU/dfba-ode-framework_ecoli-diauxie. The development version is hosted on https://gitlab.com/asuccurro/dfba-ode-framework and people interested in contributing can request access by contacting the corresponding author (A.S.).

## 5 Supplemental Material

Supplemental material for this article may be found at xxx

**FIGURE S1, S2, S3**, PDF file, 962 KB.

**FIGURE S4** EPS file, 2.2 MB.

**TEXT S1, TABLE S1**, PDF file, 228 KB.

**FIGURE S5**, PDF file, 406 KB.

**FIGURE S6, S7**, PDF file, 388 KB.

**FIGURE S8** PDF file, 449 KB.

## 6 Acknowledgments

A.S., D.S. and O.E. initiated and designed the project; A.S. developed the modeling framework, performed simulations and wrote the initial manuscript; all authors contributed to the interpretation of the results and to the final manuscript. The authors are grateful to Prof. Brice Enjalbert for kindly providing the experimental data used to compare with the simulation results. A.S. and O.E. are supported by the Deutsche Forschungsgemeinschaft, Cluster of Excellence on Plant Sciences CEPLAS (EXC 1028). A.S. was supported also by funding from the European Commission Seventh Framework Marie Curie Initial Training Network project ‘AccliPhot’ (grant agreement number PITN-GA-2012-316427). D.S. acknowledges funding from the U.S. Department of Energy (DE-SC0012627), the NIH (5R01DE024468, R01GM121950), the National Science Foundation (Grants 1457695, NSFOCE-BSF 1635070), the MURI Grant W911NF-12-1-0390, the Human Frontiers Science Program (RGP0020/2016), and the Boston University Interdisciplinary Biomedical Research Office. The funders had no role in study design, data collection and interpretation, or the decision to submit the work for publication.

